# Preserved functional organization of human auditory cortex in individuals missing one temporal lobe from infancy

**DOI:** 10.1101/2023.01.18.523979

**Authors:** Tamar I Regev, Benjamin Lipkin, Dana Boebinger, Alexander Paunov, Hope Kean, Sam Norman-Haignere, Evelina Fedorenko

## Abstract

Human cortical responses to natural sounds, measured with fMRI, can be approximated as the weighted sum of a small number of canonical response patterns (components), each having interpretable functional and anatomical properties. Here, we asked whether this organization is preserved in cases where only one temporal lobe is available due to early brain damage by investigating a unique family: one sibling born without a left temporal lobe, another without a right temporal lobe, and a third anatomically neurotypical. We analyzed fMRI responses to diverse natural sounds within the intact hemispheres of these individuals and compared them to 12 neurotypical participants. All siblings manifested the neurotypical auditory responses in their intact hemispheres. These results suggest that the development of the auditory cortex in each hemisphere does not depend on the existence of the other hemisphere, highlighting the redundancy and equipotentiality of the bilateral auditory system.

## Introduction

Mature neural tissue in human brains stores a lifetime of experiences, knowledge, and skills. As a result, brain damage sustained in adulthood often causes a loss of perceptual, motor, or cognitive functions. In contrast, the outcomes of brain damage sustained early in life are more variable. For example, perinatal strokes (strokes in fetuses or newborns), which occur in approximately 1 in 2000 term births (Dunbar and Kirton, 2019), can lead to severe long-term disabilities (e.g., Chabrier et al., 2011; Kirton and De Veber, 2013), but may also show no observable effects and go undetected for many years (e.g., Laumann et al., 2021; Tuckute et al., 2022).

Predicting the outcomes of perinatal lesions is of critical importance for diagnosis and treatment plans, and yet, the factors that determine the severity of perinatal lesion outcomes are not completely understood. The location of the injury appears to be important (Kirton and De Veber, 2013), presumably because brain areas vary in how replaceable they are, i.e. to what extent their functions could be performed by other brain areas. How replaceable a brain area is, likely depends on its maturation trajectory. Subcortical and brainstem structures develop earlier than cortical structures (Johnson, 2001). In the cortex, primary sensory and motor areas develop earlier than the association cortical areas (Johnson 2001), some of which continue to mature well into late childhood and adolescence (Giedd et al., 1999; Stiles and Jernigan, 2010). Indeed, one relatively late-developing function—language—exhibits a striking contrast between late damage to the left hemisphere (LH), which typically leads to linguistic function deficits (aphasia), and early damage (within the first few years of life) to the LH, which often results in normally developing linguistic functions, supported by the right hemisphere (RH) (e.g., Lenneberg, 1967; Newport et al., 2017, 2022; Asaridou et al., 2020; Tuckute et al., 2022; c.f. Beharelle et al., 2010). However, even in cases of damage to primary cortical areas, recovery of brain function has been reported (e.g., see Kiper et al., 2002; Knyazeva et al., 2002 for evidence of visual function recovery following extensive early damage to primary visual cortex). As such, many questions remain about the relationship between early brain damage and its long-term effects on brain function and cortical organization.

Our investigation concerns the organization of auditory cortex following extensive early unilateral lesions of the temporal lobe. Unilateral damage to temporal lobe structures may lead to severe auditory impairments when it occurs in adults (Bamiou, 2015) and children (Berticelli et al., 2021) even as early as 13 months of age (Murphy et al., 2017). However, to our knowledge, the effects of perinatal temporal lobe lesions on auditory cortical organization have not been extensively investigated. Does the auditory cortex in the intact hemisphere look typical-like, or does the lack of the contralateral homotopic areas alter its functional architecture in some way? Answers to these questions can inform our understanding of the constraints on brain development and on the possible architectures for perceptual and cognitive functions, especially those that are lateralized in typical brains.

We investigated the functional organization of auditory cortex in three members of an unusual family. Two sisters in this family (ages 54 and 55 at testing) had extensive unilateral lesions that encompassed most of the temporal lobe, likely due to perinatal stroke: in one sister, the left temporal lobe was affected (**Figure 1A**), and in the other— the right temporal lobe (**Figure 1B**). A third sister (age 53 at testing) had an intact brain (**Figure 1C**) and therefore served as a control. The affected individuals reported normal auditory, linguistic, and general cognitive abilities, as was confirmed by our behavioral testing (see <u>Supplementary Information</u> Section 1). One of the affected individuals, described in Tuckute et al. (2022) and Li et al. (2021), was not even aware of her lesion until approximately the age of 25.

**Figure 1.**
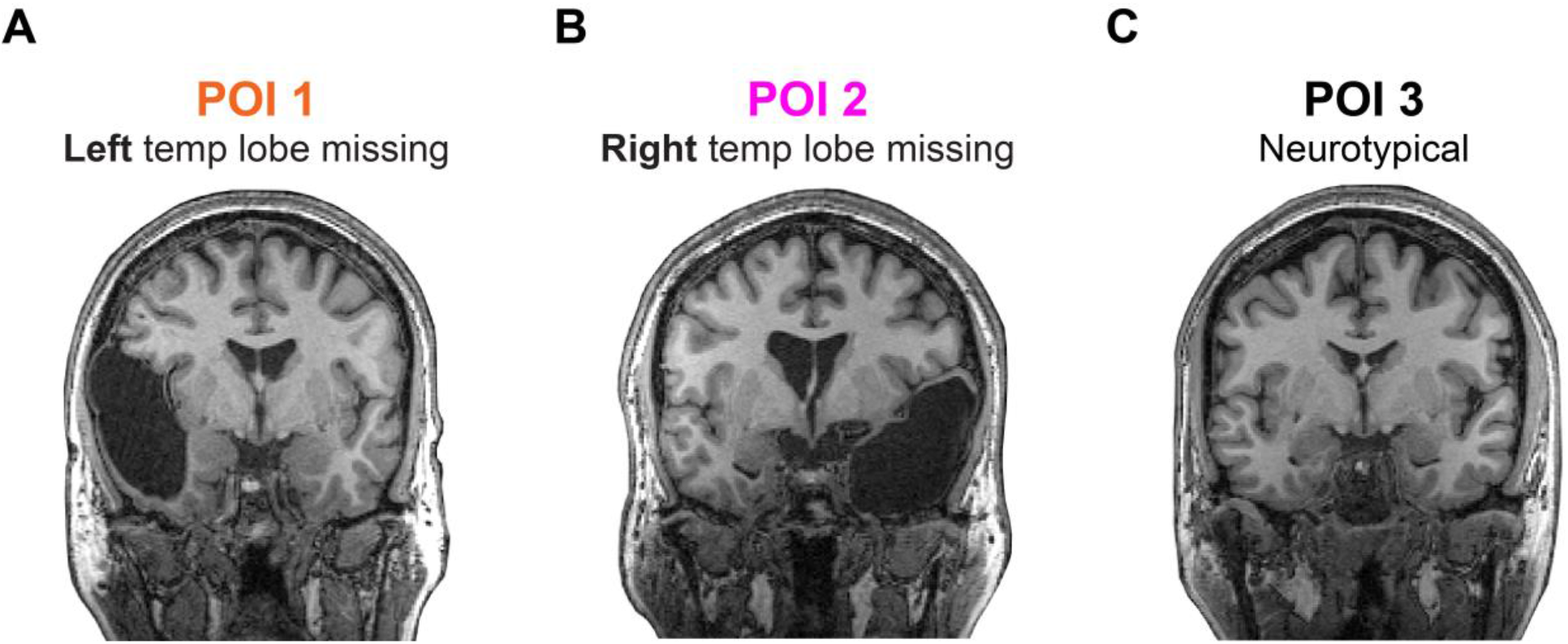
Anatomical MRI images for the three participants of interest (POI). **A**. POI 1 is missing most of the left temporal lobe from infancy; **B**. POI 2 is missing most of the right temporal lobe from infancy; and **C**. POI 3 has a typical brain.

What might we expect regarding the functional architecture of the auditory cortex in the absence of the contralateral temporal lobe? One possibility is an overall increase in auditory representation to compensate for the loss of contralateral tissue. Following brain injury, neural tissue that is able to take over the lost functions often expands (e.g., Kaas, 1991; Irvine et al., 2000). This increased representation could apply to all auditory functions (**Figure 2, Hypothesis 1**), or it could be restricted to—or especially pronounced for—auditory areas that support specific functions that lateralize to the affected hemisphere (**Figure 2, Hypothesis 2**).

**Figure 2.**
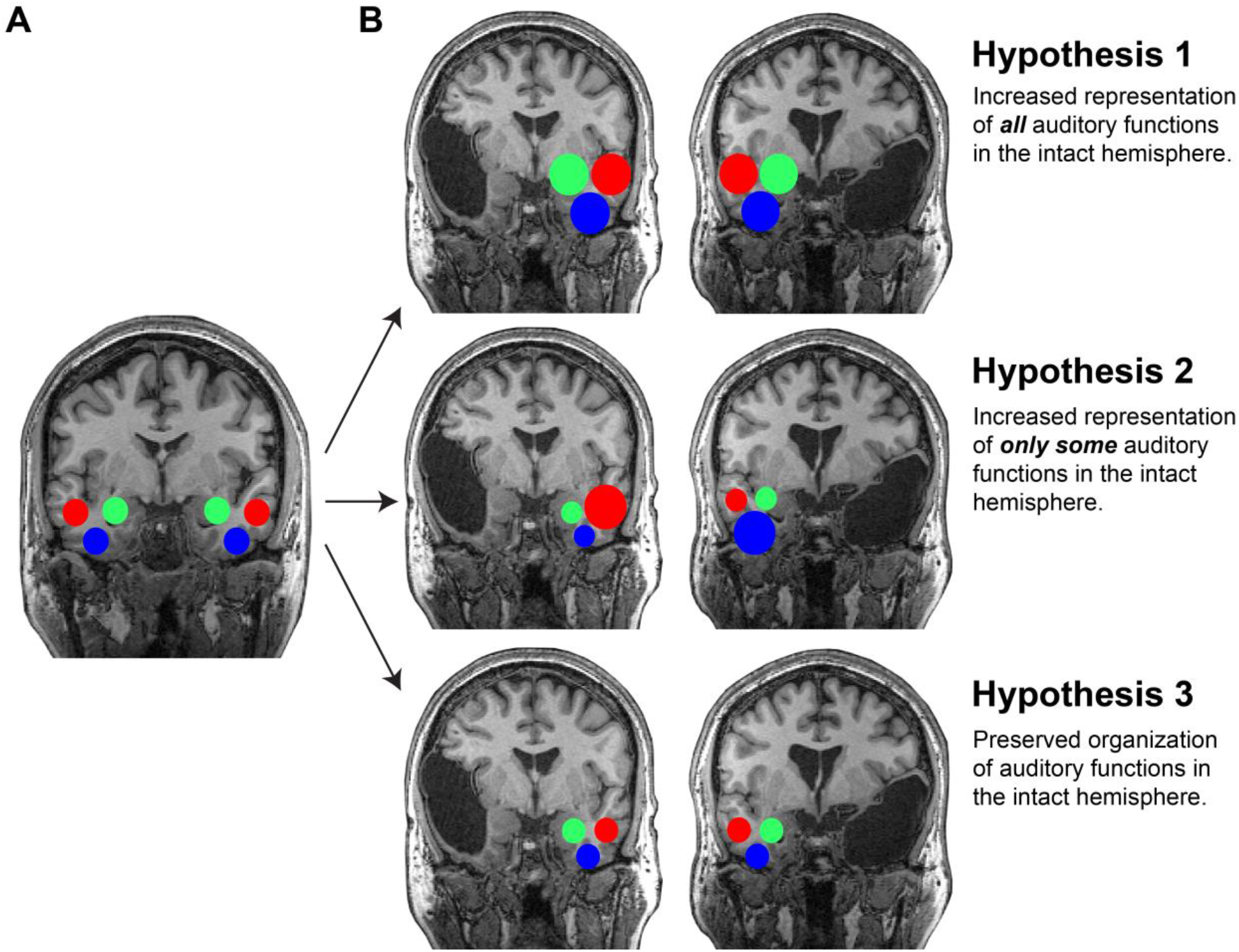
Hypotheses for functional organization of the auditory cortex in the presence of extensive contralateral temporal lobe lesions. The colored circles represent neural activation to functional auditory component (each color corresponds to a different component). **A** – Activations observed in neurotypicals. **B** – Hypotheses for activations observe in individuals missing one temporal lobe from birth.

Whether different aspects of audition preferentially depend on one or the other hemisphere is a question that has garnered much attention over the years (e.g., Zatorre et al., 2002; Poeppel, 2003; Tervaniemi and Hugdahl, 2003) and remains debated. According to one prominent proposal, the left auditory cortex is better suited for processing fast-changing auditory information, which may be important for processing the phonetic content of speech (Efron, 1963; Kimura, 1967; Schwartz and Tallal, 1980; Schonwiesner et al., 2005; Obleser et al., 2008; Albouy et al., 2020; cf. McGettigan and Scott, 2012 for arguments against this claim). The right auditory cortex, on the other hand, is postulated to be better suited for processing fine spectral modulations over longer timescales (e.g., Belin et al., 1998; Boemio et al., 2005; Abrams et al., 2011; Poelmans et al., 2012), which may be important for processing prosodic features of speech (intonation) or pitch information more broadly, including in music (e.g., Peretz, 1990; Zatorre et al., 1992, 2002; Liégeois-Chauvel et al., 1998). If the left auditory cortex is indeed more suitable for speech perception, then we might expect speech processing to take up more cortical tissue and/or elicit stronger responses in the RH in the absence of the left temporal lobe compared to neurotypicals, because the RH auditory cortex is just not as well designed for this function. And similarly, if the right auditory cortex is more suitable for music perception, then we might expect music perception to take up more cortical tissue and/or elicit stronger responses in the LH in the absence of the right temporal lobe. Alternatively, if auditory functions are redundantly supported by the two hemispheres, then we might expect the auditory organization in the intact hemisphere to look typical-like (**Figure 2, Hypothesis 3**).

To investigate the organization of the auditory cortex in our participants of interest (POI), we adopted a paradigm developed by Norman-Haignere et al. (2015). Norman-Haignere and colleagues recorded fMRI responses to a diverse set of natural sounds and, using a data-driven approach, uncovered six response components that capture most of the explainable variance in auditory cortical responses (see Boebinger et al., 2021 for replication). Component 1 and 2 were selective for low and high frequency acoustic information, respectively, as confirmed by a separate assessment of tonotopy (Norman-Haignere et al., 2015); Components 3 and 4 were selective for spectrotemporal modulations that tend to be present in environmental sounds and pitched sounds, respectively; and Components 5 and 6 were highly selective for speech and music, respectively (**Figure 3D**). We tested whether this auditory cortical organization is preserved when the auditory cortex in the contralateral hemisphere is missing from birth/infancy.

**Figure 3.**
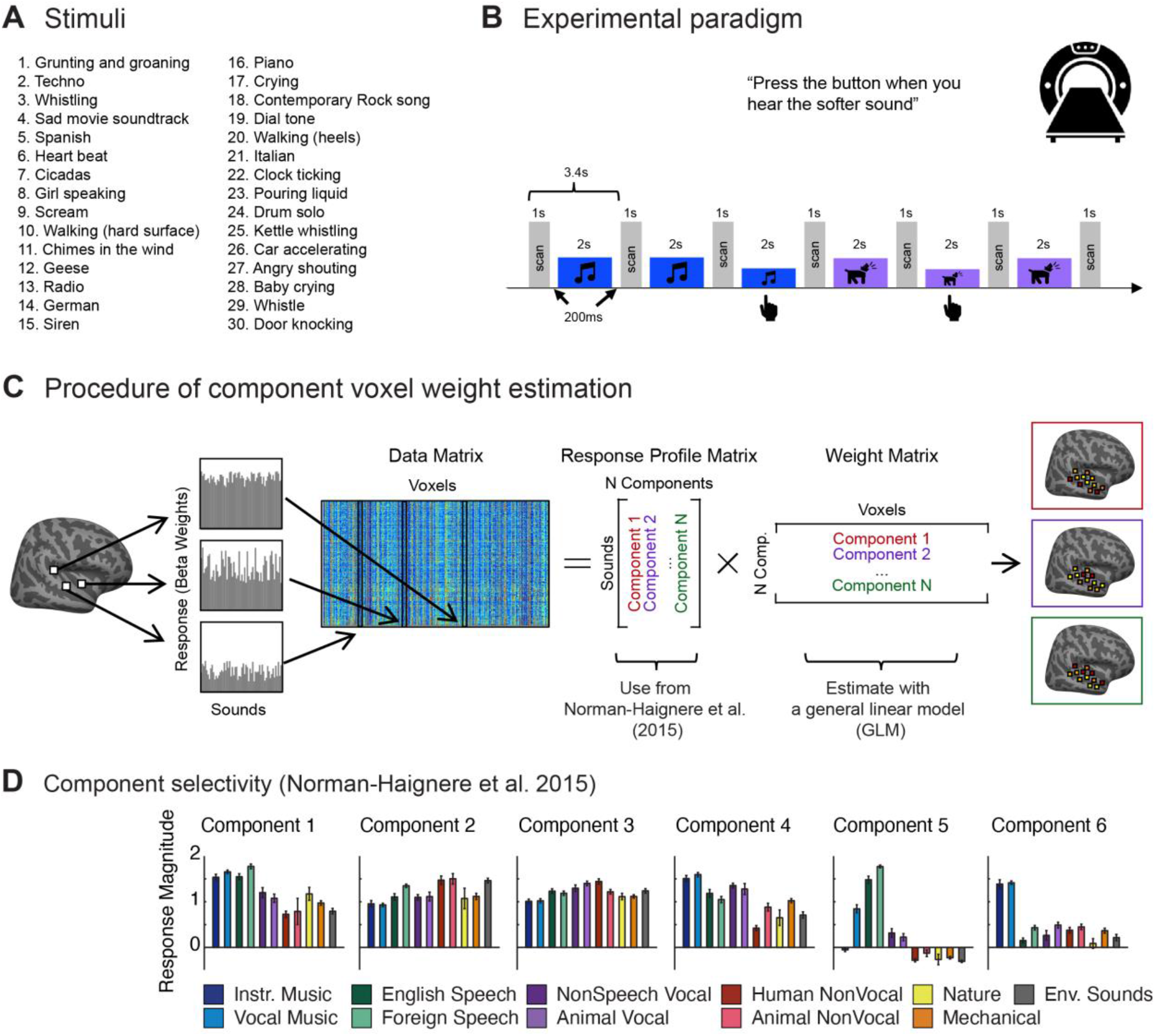
Experiment design and auditory component estimation procedure. Adapted from Norman-Haignere et al. 2015 and Boebinger et al. 2021 with permission. **A** – The 30 sound stimuli presented in the experiment. This set of 30 stimuli was chosen from the original set of 165 natural sounds in order to optimize the detection of the 6 components (Boebinger et al 2021, <u>Methods</u>). The 30 sounds are ordered here by a score of how well they were suited for optimizing the detection of the 6 components. **B** – Experimental paradigm. Each 2-s sound stimulus was repeated three times consecutively, with one repetition (the second or third) being 8 dB quieter. Subjects were instructed to press a button when they detected this quieter sound. One fMRI volume was acquired in the silent period between stimuli (sparse scanning). **C** – Procedure of component voxel weight estimation. Whereas Norman-Haignere et al. 2015 and Boebinger et al. 2021 estimated both a component response profile matrix and a weight matrix, we used the component response matrix from Norman-Haignere et al. and estimated just the voxel weight matrix given our data matrix, using a general linear model (GLM). **D** – Component responses averaged across sounds from the same category. (Sounds were assigned to categories by an independent set of participants in an online study, as described in Norman-Haignere et al., 2015.) Error bars represent one standard error of the mean across sounds from a category, computed using bootstrapping (10,000 samples).

## Results

We measured fMRI responses to a set of 30 natural sounds from a variety of categories (**Figure 3A**) in three participants of interest (POIs): three sisters, with one missing most of her left temporal lobe (POI 1), one missing most of her right temporal lobe (POI 2), and the third neurotypical with both temporal lobes intact (POI 3), as well as in a control population of 12 participants. The 30 sounds presented in this experiment were a subset of the 165 sounds originally tested in Norman-Haignere et el. (2015) and were selected to optimally identify component weights (Boebinger et al. 2021, <u>Methods</u>).

We then projected the response of each voxel onto the six auditory response components from Norman-Haignere et al. (2015) using a general linear model (GLM) with ordinary least squares regression (**Figure 3C, D**). This projection provided estimated weights for each of the six components for each voxel and participant. We studied the component responses (weights) within the auditory cortex, which was defined as the conjunction of 15 anatomical parcels (Glasser et al. 2016; selection of parcels as in Boebinger et al. 2021; <u>Methods</u>; **Figure 4**).

**Figure 4.**
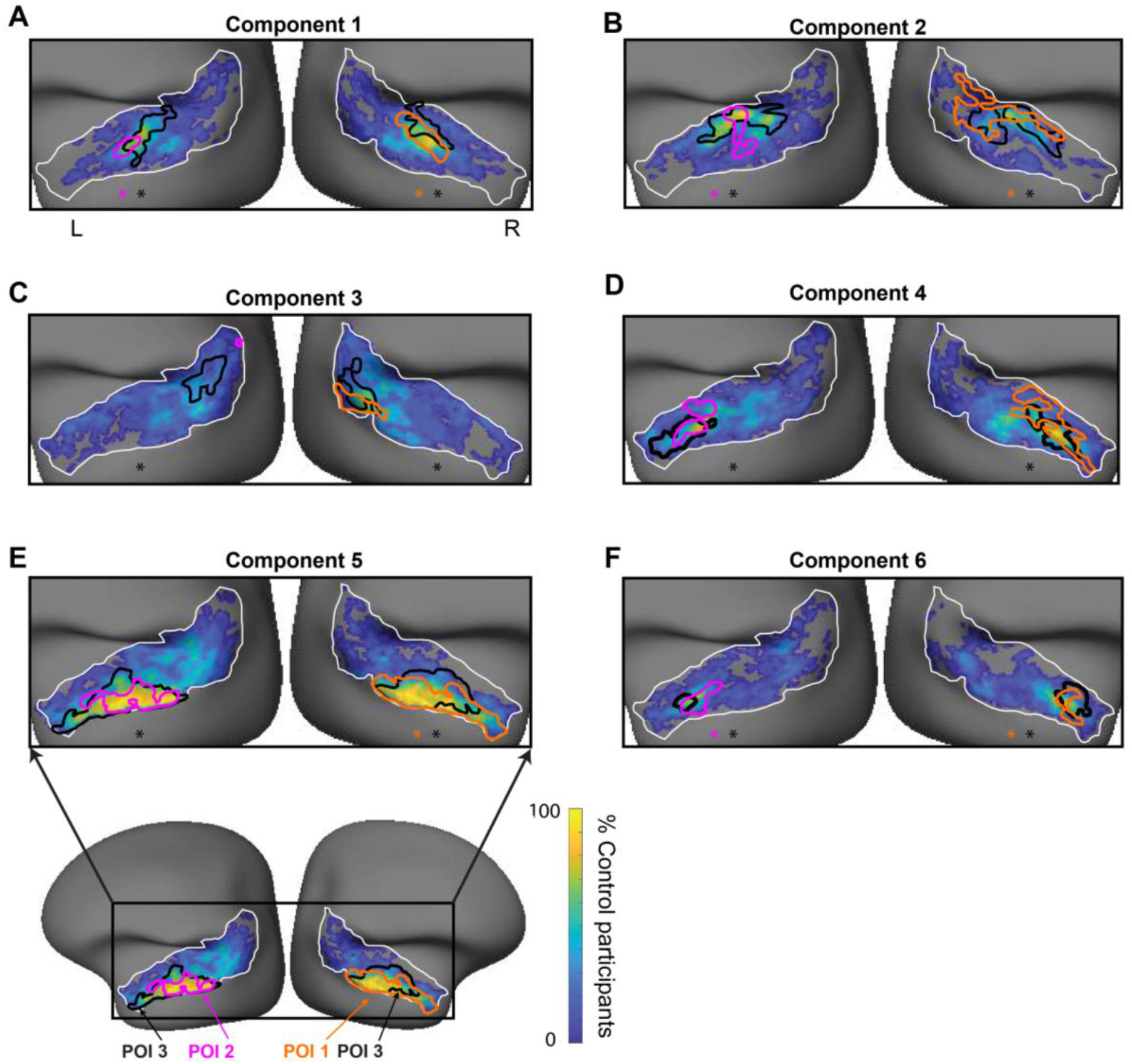
Functional auditory cortex components in intact hemispheres of the three participants of interest, relative to probability maps derived from the 12 control participants. For each component and hemisphere, probabilistic maps (blue to yellow color scale) depict for each voxel the percentage of control participants who show a significant response (p<0.05). Voxels for which no control participant showed a significant response were left gray, as were voxels outside of the union of the 15 parcels chosen to comprise auditory cortex (depicted by the white outline, Glasser et al., 2016, selection of parcels as in Boebinger et al 2021, <u>Methods</u>). On top of these maps, the largest contiguous cluster of voxels passing the same threshold is outlined in orange for POI 1, pink for POI 2, and black for POI 3. Stars indicate a significant (p<0.05) Pearson correlation across the 15 anatomical parcels between the spatial pattern of each POI’s component response and the average component response of the 12 neurotypical participants. See all maps at <u>osf.io/qrx5n/.</u>

Before performing further analyses, we calculated the proportion of the variance explained by the six components. For each voxel in the auditory cortex of each participant, we estimated the amount of variance that was explained by the six components relative to the total variance in that voxel: 1-Var(ŷ-y)/Var(y) where ŷ is the estimate of the signal using all six components and y is the observed response (ŷ + residual of the model). For the 12 neurotypical participants, the median voxel variance explained (across the two hemispheres) ranged from 29.9% to 40.3% (note that these values are expected to be well below 100% because we examine variance that is not corrected for the noise in the data). The median voxel variance explained in each of the 3 POIs fell right in the middle of this range (POI 1: 35.7%, POI 2: 33.3%, and POI 3: 36.7%), which suggests that the POIs are comparable to the controls in terms of how well their auditory neural activity is captured by the six components.

All subsequent analyses were performed component-wise within the auditory cortex of the intact hemisphere in the POIs, and the results were compared to the same hemisphere in the neurotypical controls using statistical methods designed for comparing single cases to normative distributions (Crawford test, Crawford & Howell (1998); <u>Methods</u>). To compare the organization of auditory cortex in our POIs to that of neurotypical controls, we examined three functional characteristics of the observed responses: 1) the response magnitude and spatial extent of each component; 2) the spatial layout of the components; and 3) the reliability of the component topographies across scanning runs.

### 1) Response Magnitude and Spatial Extent

For each of the three POIs and each of the 12 control participants, the mean response magnitude (weights) and the spatial extent (number of significant voxels at the p<0.05 level within the auditory mask) were calculated for each component. The mean response magnitudes and spatial extents of all components for the three POIs fell within the range of the distribution of the 12 controls. No significant differences were observed between the mean response magnitudes (**Figure 5A**) or the significant voxel counts (**Figure 5B**) of any of the POIs and the control participants, for any of the six components (p>0.05; Crawford test, FDR corrected for the 6 components, **Table S1**).

**Figure 5.**
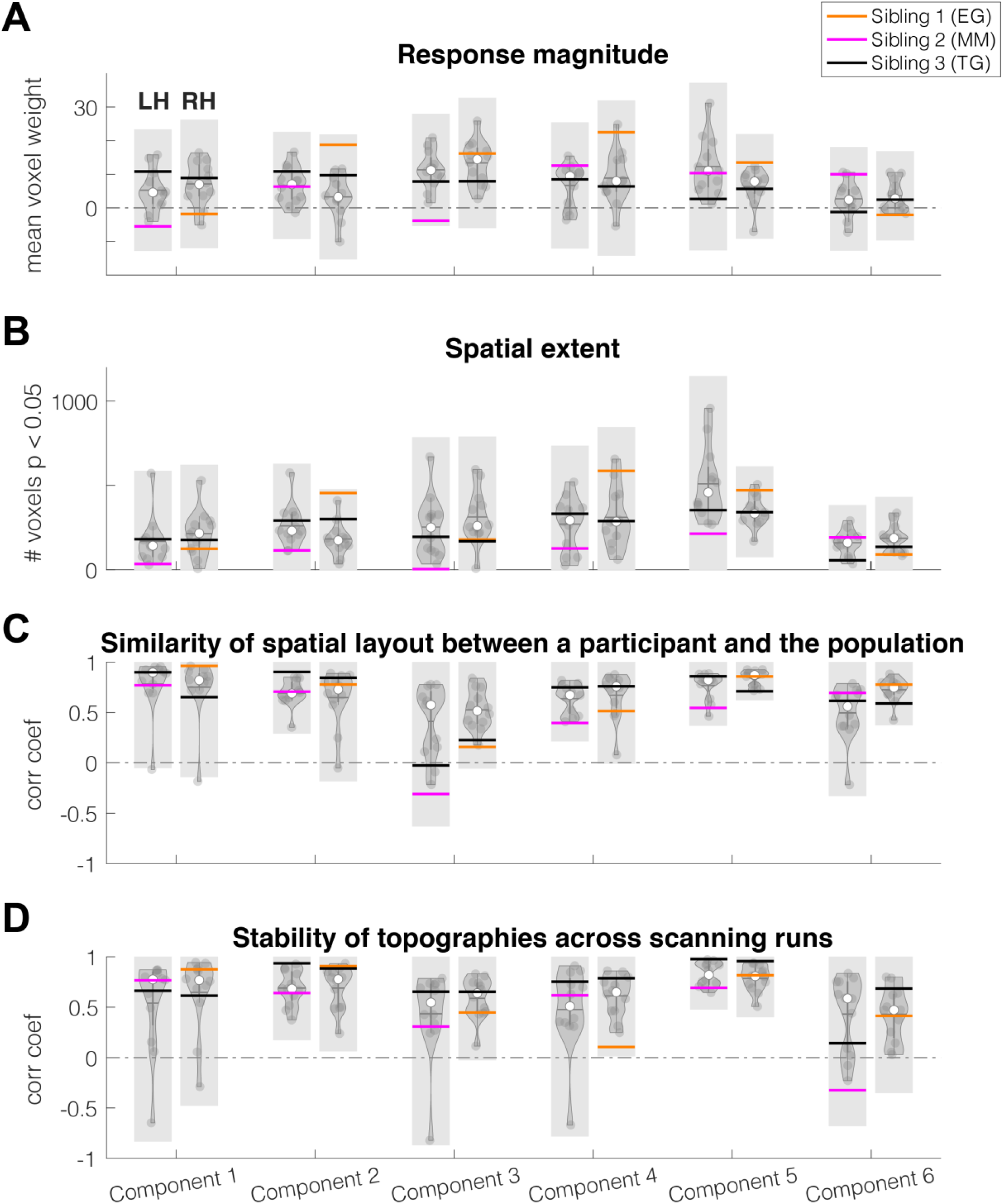
Comparison—for the different properties of the six components—between the three participants of interest and the 12 control participants. Each violin plot represents the 12 control participants. For each property (panel, A-D) and component (labels on the lowest x-axis), the two violins represent the left hemisphere (left) and right hemisphere (right). Response magnitude (A) and Spatial extent (B) are proxies for the amount of neural activity, whereas Similarity of spatial layout between a participant and the population (C) and Stability of topographies across scanning runs (D) reflect information about the robustness of the fine-grained activation patterns. See <u>Results</u> and <u>Methods</u> for more details. Horizontal lines represent POI 1 (orange) missing most of the left temporal lobe from infancy, POI 2 (magenta) missing most of the right temporal lobe from infancy, and POI 3 (black) with a typical brain. Shaded regions reflect the ‘typical’ range which the POIs would need to fall outside of to significantly deviate from the control population according to the Crawford test (Crawford & Howell (1998), <u>Methods</u>). In other words, values within this range would not significantly deviate (p<0.05) from the null Crawford distribution, parameterized by the control participants, whereas values outside this range would indicate significant deviation. The bounds of this range were calculated as the test values to this distribution that would result in p=0.05.

### 2) Spatial Layout

We examined the anatomical spatial distribution of response components in two ways. First, we plotted each component for each POI on top of a probabilistic map which we computed for the 12 control participants using the binarized significant (p<0.05, uncorrected) voxel responses. This allowed for a visual assessment of each component in the POIs (binarized in a similar way) relative to the control population. The POIs’ components fell qualitatively within the boundaries of the neurotypical component distributions (**Figure 4**). The only exception was Component 3 (environmental sounds) in the (left) auditory cortex of POI 2, which was almost absent (**Figure 4C**). However, note that the location of Component 3 was more variable than the other components across the 12 neurotypical control participants (**Figure 4C**, also see Figure 2A in Boebinger et al. 2021 and Figure S5 from Norman-Haignere et al. 2015, demonstrating the lower reliability of Component 3 compared to other components between and within participants, respectively).

Second, we quantified the similarity of the overall spatial component layout in the POIs vs. controls via correlations across 15 anatomical parcels chosen to comprise auditory cortex (Glasser et al. 2016; selection of parcels as in Boebinger et al. 2021; <u>Methods</u>). We chose to project the responses to this lower-dimensional space because establishing voxel-wise functional correspondences across individuals is challenging due to inter-individual variability in the auditory cortex (e.g., Moerel et al., 2014; Ren et al., 2021). In particular, for each component, a Pearson correlation was computed across the 15 parcels between each POI’s component response and the average component response of the controls. POI 1 presented with a significant correlation to the control group for four of the six components (Components 1,2,5,6, p<0.01 FDR corrected, **Figure 4, Table S2**), POI 2—for three of the six components (Components 1,2,6), and POI 3—for five of the six components for each of the two hemispheres (Components 1,2,4,5,6). Component 3 was not significantly correlated with the control group in any of the POIs.

To investigate the extent to which the observed correlation coefficients were within the expected range of individual differences, we performed the same correlation analysis reported above for each of the 12 control participants via their correlation with the average of the remaining 11 control participants. We then tested whether the values of the correlation coefficients obtained for the POIs (comparing the POIs to the average of the 12 controls) differed significantly from the values of the correlation coefficients obtained for the control group (comparing each control participant to the average of the remaining 11 controls). No significant differences were observed for any of the components between a) the correlation coefficients obtained when comparing each of the POIs to the control group and b) the correlation coefficients obtained when comparing each of the 12 control participants to the average of the remaining 11 controls (all p<0.01, Crawford test, **Figure 5C**; **Table S1**).

### 3) Reliability of the Component Topographies Across Scanning Runs

To examine the stability of the component topographies over time, we correlated the response profiles—across the same 15 parcels (Glasser et al., 2016) that were used in the analysis of the spatial layout—between odd- and even-numbered scanning runs. POI 1 showed significantly stable patterns of activation for 3 of the 6 components (Components 1, 2, 5), POI 2—for 4 or the 6 components (Components 1, 2, 4, 5), and POI 3 for 5 of the 6 (components 1, 2, 3, 4, 5) LH components and for all RH components (**Table S2**). As in the previous section, this process was repeated for each of the control participants to establish a null distribution of between-run correlations. No significant differences were observed for any of the components between a) the correlation coefficients obtained for the POIs across scanning runs and b) the correlation coefficients obtained for each of the 12 controls across scanning runs (**Figure 5C**; **Table S1**).

## Discussion

We examined a rare case of two siblings who are each lacking (most of) one of their temporal lobes from infancy: one in the left hemisphere, the other in the right. We asked whether the organization of their auditory cortex in the intact hemisphere is similar to that observed in typical brains. Using fMRI, we measured neural responses to natural sounds in these two siblings, a third (neurotypical) sibling, and 12 additional neurotypical control participants and decomposed those responses into six functional components (Norman-Haignere et al. 2015, Boebinger et al. 2021). Our findings suggest that auditory cortical organization is preserved in the intact hemispheres of the sisters lacking one temporal lobe from infancy.

In particular, i) all components discovered by Norman-Haignere et al. were reliably detectable in each participant of interest; and ii) their anatomical locations were similar to what is observed in typical brains. Furthermore, using statistical methods designed for comparing single cases to normative distributions, we searched for significant differences between the participants of interest and the control group in several component properties, including response (weight) magnitude, spatial extent, overall spatial layout, and stability of the activation patterns over time (across scanning runs). At least based on the current (relatively small) control sample, iii) we observed no significant differences between the POIs and the controls. Although the lack of significant differences between the participants of interest and the controls could be construed as a null finding, the robust presence of all the components and their similar spatial distribution are positive results that suggest preserved functional organization in spite of extensive early brain damage.

The original findings of Norman-Haignere et al. (2015) as well as their replication in Boebinger et al. (2021) did not manifest strong hemispheric asymmetries: all six components were roughly symmetric in their response profiles and spatial extent, with no significant hemispheric differences in the average weight for any of the components. Even the speech component was similarly robust in the left and right hemispheres, in contrast with some claims of left-lateralized speech responses in infancy (e.g., Dehaene-Lambertz et al., 2002; Peña et al., 2003; Vannasing et al., 2016; Daneshvarfard et al., 2019; cf. Cristia et al., 2014). Our findings further extend this lack of asymmetry in demonstrating independence of each hemisphere’s auditory cortex from the existence of the contralateral one. The speech component in POI 1’s right hemisphere is similar to the speech component in the RH of neurotypical individuals. Our findings suggest that speech processing can successfully rely on the RH auditory cortex, and more generally, that auditory functions can be supported by auditory cortex in just one hemisphere, either the left or the right one.

The finding that the RH auditory cortex can support speech perception aligns with reports of high-level language processing being successfully supported by the right hemisphere in some individuals with early left-hemisphere damage (Newport et al., 2022; Tuckute et al., 2022; see François et al., 2021 for a review). Such individuals develop linguistic abilities at the same time as their neurotypical peers and exhibit no language difficulties as adults. If the (earlier-developing) speech perception abilities could *not* be supported by the right hemisphere (or at least not as well as by the left hemisphere), individuals with early left-hemisphere damage should inevitably exhibit delays in language acquisition and/or lasting deficits in language processing.

The apparent equipotentiality of the two hemispheres for auditory processing suggests that any anatomical asymmetries in auditory cortex are not necessary for normal cortical functioning. In particular, anatomical asymmetries have been reported in fetal and infant brains in auditory cortical areas, including Heschl’s gyrus and Planum temporale (e.g., Geschwind and Levitsky, 1968; Chi et al., 1977; Wada, 2011), as well as brain areas / tracts surrounding the auditory cortex (e.g., the arcuate fasciculus; Dubois et al., 2009; superior temporal sulcus; Glasel et al., 2011; Leroy et al., 2015). However, given that auditory processing can be supported by either hemisphere, these asymmetries do not appear to be critical for auditory function.

Overall then, our results demonstrate that neurotypical-like organization of the auditory cortex emerges in the absence of the contralateral temporal lobe (regardless of which hemisphere is affected), highlighting the independence and equipotentiality of the auditory cortex of the two hemispheres and the redundancy in the auditory cortical system in humans.

## Methods

### Participants of Interest (POIs)

The first POI, henceforth referred to as POI 1 contacted professors at MIT Brain and Cognitive Sciences (BCS) in February 2016 volunteering to participate in studies of her brain, which she reported had no left temporal lobe. POI1 had never suffered any head traumas or injuries, but discovered this feature when an MRI scan was performed in 1987 when she was 25 years old during treatment for depression. Additional scans were performed in 1988, 1998, and 2013 without any reported changes. Despite this supposedly congenital condition, POI1 is highly educated, with an advanced professional degree, and reports no problems with vision, except nearsightedness (corrected with glasses), and had even studied Russian as a foreign language in adulthood, achieving high proficiency. POI1 (right-handed, 54-years old at testing) participated in behavioral and fMRI testing at MIT in October 2016.

The second POI, henceforth referred to as POI 2 is POI 1’s sister. During testing for some vision problems in 1981 when she was 17 years old, POI 2 discovered that she had no right temporal lobe. POI 2 (right-handed, 55-years-old at testing) participated in behavioral and fMRI testing at MIT in September 2019.

The third POI, henceforth referred to as POI 3 is the third sister of POI 1 and POI 2. POI 3 has a neurotypical brain and serves as a close control to the sisters, discounting potential sources of variation from the control population outlined below. POI 3 (right-handed, 57-years-old at testing) participated in behavioral and fMRI testing at MIT in September 2019.

### Control Participants

In addition to the three sisters, 12 neurotypical participants (mean age = 27.8, std = 4.1 years, 3 females, 11 right-handed and 1 ambidextrous, according to self-report) were recruited for fMRI testing at BCS. These participants were recruited from MIT and the surrounding Cambridge/Boston, MA community and were paid for their participation. All participants had normal hearing and vision.

These data were originally collected for the purpose of other studies and were re-analyzed here. The protocol for these studies was approved by MIT’s Committee on the Use of Humans as Experimental Subjects (COUHES). All participants gave written informed consent in accordance with the requirements of this protocol.

### Stimuli & fMRI Task Design

An abridged version of the task used in Norman-Haignere et al. (2015) was used to evoke responses from participants to a selection of 30 2-second natural sounds (included in supplemental; <u>osf.io/qrx5n/</u>). These 30 sounds were chosen as the subset of the 165 sounds that were best able to identify the 6 components. This was done using a greedy algorithm to search for subsets of sounds that had a high but uncorrelated response variance across the components (Boebinger et al 2021).

Stimuli were presented during scanning in a “mini-block design,” in which each 2-second sound was repeated three times in a row. During scanning, stimuli were presented over MR-compatible earphones (Sensimetrics S14) at 75 dB SPL. Each stimulus was presented in silence, with a single fMRI volume collected between each repetition (i.e. “sparse scanning”; Hall et al., 1999). To encourage participants to pay attention to the sounds, either the second or third repetition in each “mini-block” was 8dB quieter (presented at 67 dB SPL), and participants were instructed to press a button when they heard this quieter sound. All sounds were presented diotically and thus we do not examine hemispheric differences in spatial coding.

Each run of the experiment included all 30 stimuli and lasted approximately 6.5 minutes. The POIs each completed 6 runs, and the control participants completed 6-10 runs. To confirm initial data quality, motion was quantified for each run of each participant. The mean RMS deviation of all runs across all participants was <1mm (0.18±0.11mm for neurotypical participants; 0.22±0.14mm for POIs), and the run-level motion parameters of the POIs did not differ from the neurotypical participants (two-sample t-test with unequal variance; t=1.25; two-sided p=0.22).

### fMRI Data Acquisition

Structural and functional data were collected on a 3 Tesla, 32-channel head coil, Siemens Trio scanner at the Athinoula A. Martinos Imaging Center at the McGovern Institute for Brain Research at MIT. T1-weighted structural images were collected via 176 sagittal slices with 1mm isotropic voxels (TR=2530ms, TE=3.48ms). Functional BOLD data were acquired in 31 4mm thick near-axial slices acquired in the interleaved order (with 10% distance factor) using an EPI sequence with the following parameters: 90 degree flip angle, GRAPPA acceleration factor 2, 2.1mm×2.1mm in-plane resolution, field of view of in the phase encoding (A>P) direction 200mm and matrix size 96mm×96mm, TR=2000ms and TE=30ms. Prospective acquisition correction (Thesen et al., 2000) was used to adjust gradient position based on the participant’s motion from the previous TR. The first 10s of each run were excluded to allow for steady state magnetization.

### fMRI Preprocessing

Anatomical and functional data were preprocessed using FreeSurfer (https://surfer.nmr.mgh.harvard.edu/), FSL (https://fsl.fmrib.ox.ac.uk/fsl/fslwiki/FSL), and supporting custom MATLAB scripts. Surface reconstructions were generated using FreeSurfer recon-all (Dale et al, 1999) and the reconstructions of the lesions were confirmed visually by the authors. Motion correction was implemented using FSL MCFLIRT (Jenkinson et al., 2002), and functional images were skull-stripped using FSL BET2 (Jenkinson et al., 2005). Functional images were then coarsely registered to their high-resolution anatomical counterparts using FSL FLIRT (Jenkinson et al., 2001) and fine-tuned using a boundary-based alignment algorithm referred to as BBRegister (Greve & Fischl, 2009). Registrations were confirmed manually by the authors, especially in the most susceptible regions around the lesions of the sisters. Following registration, functional data were resampled from 3D volume space to 2D cortical surfaces using FreeSurfer mri_vol2surf, and aligned to the FsAverage template brain using FLIRT and BBRegister. Finally, data were smoothed using a 3mm FWHM Gaussian kernel and downsampled to a 1.5×1.5mm flattened surface grid in MATLAB.

### fMRI First-Level Modeling

Effects in each vertex of the surface grid were estimated using a General Linear Model (GLM). There was a separate regressor for each component. This regressor was computed by creating a boxcar function for each stimulus whose height was equal to the component’s response to that stimulus, as estimated in our prior studies. These boxcar functions were then convolved with the canonical hemodynamic response function (HRF). Solving this GLM calculated a beta weight for each component of each voxel and these estimates formed the basis of subsequent analyses. Since fMRI data is neither independent across time nor Gaussian distributed, the significance of these beta estimates was evaluated using a permutation test. For each voxel, a null distribution was estimated by randomly permuting the order of stimuli in the paradigm file, and re-running the analysis (n=1000). The above process was originally performed using all runs, and was later repeated using only odd/even runs for subsequent stability analyses.

### Definition of Stimulus Component Weights

The natural sound component weights used in this study were defined via Norman-Haignere et al using a hypothesis-free voxel decomposition of auditory cortex responses to a large collection of natural sounds (superset of sounds presented in this study). In voxel decomposition, voxels are approximated as a weighted sum of a small number of canonical response patterns. This approximation problem is ill-posed and must be constrained by additional statistical criteria, which was accomplished by searching for components that have a non-Gaussian weight distribution across voxels (see Norman-Haignere et al., 2015 for a detailed discussion). Once the component response patterns are known, the weights can be estimated using ordering least-squares regression, as described above, which requires much less data than computing the full decomposition (which depends upon higher-order statistics such as the skew and kurtosis which require large amounts of data to robustly measure). In the original dataset, such a decomposition yielded a set of 6 components, which explained >80% of the noise-corrected variance. These six components have since been replicated twice in two independent populations (Boebinger, et al (2021). We used the response patterns from the six components from Norman-Haignere et al., (2015) to calculate the component weights reported here.

### Definition of Anatomical Auditory ROIs

All analyses were restricted to an anatomically defined region of the broader auditory cortex, as in Boebinger et al (2021). This region was defined using the Glasser et al. (2016) atlas based on a multi-modal parcellation of the human cerebral cortex, with the goal of broadly encompassing sound-responsive cortex. The parcels for the following regions were resampled to the surface grid space used in this study, and used to constrain the ROI for analysis: Primary Auditory Cortex, PeriSylvian Language Area, Superior Temporal Visual Area, Area 52, RetroInsular Cortex, Area PFcm, Area TA2, Area STGa, ParaBelt Complex, Auditory 5 Complex, Area PF Complex, Medial Belt Complex, Lateral Belt Complex, Auditory 4 Complex, and Para-Insular Area.

### Critical Analyses Note on lesions

All subsequent analyses were performed using only non-lesioned hemisphere data, constraining only RH for POI 1 and only LH for POI 2. Thus all analyses of RH activity would evaluate POI 1 RH and POI 3 RH in relation to neurotypical RH, and all analyses of LH activity would evaluate POI 2 LH and POI 3 LH in relation to neurotypical LH.

### Note on statistical testing

In order to compare single subject data to a null group, a special statistical test, developed by Crawford & Howell (1998), was used in several of the following analyses. The Crawford test is a Bayesian statistical test designed to evaluate the typicality of a single value against a set of control values. It behaves similarly to frequentist methods, e.g. fitting a Gaussian and performing a one-sample z-test; however, rather than deriving *population parameters* from the control sample, the Crawford test treats the control statistics as *sample statistics*, preventing inflation to the Type I error rate in cases where the control sample is modest in size, e.g. *N* around 10 (Crawford and Garthwaite, 2007). In this study, it was used to compare values extracted from POI 1’s and POI 3’s RH to a set of values from the neurotypical RH, and values extracted from POI 2’s and POI 3’s LH to a set of values from the neurotypical LH. All reported Crawford p-values are two-tailed.

### Probabilistic map

For each of the three POIs and each of the 12 neurotypical control participants, the significant voxels (p<0.05, uncorrected) responsive to each component within the auditory cortex ROI were extracted, and assigned a value of 1 if significant, and 0 otherwise. From these neurotypical-participant-level binary maps, the probabilistic map of the response profile was generated by calculating the mean of the binary maps. For example, if a voxel was significant in 6 of 12 participants, it’s value in the probabilistic map would be 0.5. Each of the POIs responses (binarized in the same way) were plotted onto this atlas to qualitatively identify a spatial correspondence in activity. For visualization purposed we plotted just the largest contiguous cluster of the binarized POIs response map (**Figure 4**).

### Response Magnitude and Spatial Extent

For each of the three POIs and each of the 12 neurotypical control participants, the mean effect size (beta; response magnitude) and the count of significant voxels (p<0.05; spatial extent) of each component were calculated within the auditory parcel of each hemisphere. Using a Crawford test, each of the POIs’ statistics were compared to the neurotypical group. Results were FDR-corrected (Benjamini and Hochberg, 1995) for the number of components (n=6).

### Spatial Layout

For each component, the spatial response of each participant within the auditory ROI was extracted as a vector of the mean effect sizes within the 15 auditory parcels (Glasser et al., 2016). Then, for each POI, Pearson’s linear correlation coefficient was calculated between the POI’s spatial component response, and the mean neurotypical spatial component response. These correlation coefficients were evaluated analytically for significance (alpha=0.05). Subsequently, this process was repeated for each of the 12 neurotypical participants via their correlation with the mean of the remaining 11 neurotypical participants. Then, for each component, the correlation coefficients between each of the POIs and the neurotypicals were compared to the correlation coefficients within the neurotypical group via a Crawford test. Both correlation and Crawford analysis results were FDR-corrected for the number of components (n=6).

### Stability of the Component Topographies Over Time

Using component activations (per parcel) calculated from only the odd or even runs of the experiment separately, the spatial correlation between these sets was calculated for each component of each participant in their relevant hemispheres using Pearson’s linear correlation coefficient. These correlations were evaluated for significance analytically (alpha=0.05) as a measure of component spatial stability. The spatial stability of each of the POIs’ responses was then compared to the neurotypical group via a Crawford test. Both correlation and Crawford analysis results were FDR-corrected for the number of components (n=6).

## Acknowledgements

We would like to acknowledge the Athinoula A. Martinos Imaging Center at the McGovern Institute for Brain Research at MIT, and its support team (Steve Shannon and Atsushi Takahashi). We thank the participants of interest who agreed to participate in our study, as well as former and current EvLab members, especially Greta Tuckute and Niharika Jhingan, for their help with fMRI data collection and analysis. TIR was supported by the Zuckerman-CHE STEM Leadership Program and by the Poitras Center for Psychiatric Disorders Research. EF was supported by NIH awards R01-DC016607, R01-DC016950, and U01-NS121471 and research funds from the the McGovern Institute for Brain Research, Brain and Cognitive Sciences department, and the Simons Center for the Social Brain.

## Supplementary Information

### 1. Behavioral assessment of language and general cognitive skills for the POIs

#### 1.1 Language assessment

To assess language skills, 4 standardized language assessment tasks were used: i) an electronic version of the Peabody Picture Vocabulary Test (PPVT-IV) (Dunn and Dunn, 2007); ii) an electronic version of the Test for Reception of Grammar (TROG-2) (Bishop, 2003); and iii) the Western Aphasia Battery-Revised (WAB-R) (Kertesz, 2006). PPVT-IV and TROG-2 target receptive vocabulary and grammar, respectively. In these tasks, the participant is shown sets of four pictures accompanied by a word (PPVT-IV, 72 trials) or sentence (TROG-2, 80 trials) and has to choose the picture that corresponds to the word/sentence by clicking on it. WAB-R (Kertesz, 2006) is a more general language assessment for persons with aphasia. It consists of 9 subscales, assessing 1) spontaneous speech, 2) auditory verbal comprehension, 3) repetition, 4) naming and word finding, 5) reading, 6) writing, 7) apraxia, 8) construction, visuospatial, and calculation tasks, and 9) writing and reading tasks.

### 1.2 General cognitive assessment

To assess general cognitive skills, 2 tasks were used: i) an electronic version of the Kaufman Brief Intelligence Test (KBIT-2) (Kaufman and Kaufman, 2004), and ii) the 3-pictures version of the Pyramids and Palm Trees Test (Howard and Patterson, 1992). The former consists of three subtests – two verbal (Verbal Knowledge and Riddles) and one non-verbal (Matrices) – and is used to assess general fluid intelligence. The Verbal Knowledge subtest consists of 60 items measuring receptive vocabulary and general information about the world; the Riddles subtest consists of 48 items measuring verbal comprehension, reasoning, and vocabulary knowledge; and the Matrices subtest consists of 46 items that involve both meaningful (people and objects) and abstract (designs and symbols) visual stimuli that require understanding of relationships among the stimuli. The Pyramids and Palm Trees test assesses non-verbal semantic cognition. The task consists of 52 trials. On each trial the participant is shown a test picture (e.g., an Egyptian pyramid) and two other pictures (e.g., a palm tree and a fur tree) and asked to choose the picture that is semantically related to the test picture (in this case, a palm tree is the correct answer). For both tests, the POI’s performance was evaluated against existing norms.

### 1.3 Behavioral results

#### 1.3.1 POI 1

In line with POI 1’s self-report, she performed within normal range on all language and general cognitive tasks. She got 90% correct on PPVT, 99% correct on TROG, and 97.6, 98.6, and 98.4 on the aphasia, language, and cortical quotients of the WAR-B (the criterion cut-off score for diagnosis of aphasia is an aphasia quotient of 93.8). POI 1’s performance was therefore not distinguishable from the performance of neurotypical controls. Her KBIT scores were 130 (98th percentile) on the verbal composite assessment (across the two subtasks), 54 (79th percentile) on the non-verbal assessment, and 122 (93rd percentile) overall composite assessment. She answered 51 of the 52 questions correct on the Pyramids and Palm Trees task.

#### 1.3.2 POI 2

POI 2 performed within normal range on all language and general cognitive tasks. She got 83.3% correct on PPVT, 96.25% correct on TROG, and 100 and 100 on the aphasia, and language quotients (cortical quotients was not completed) of the WAR-B (the criterion cut-off score for diagnosis of aphasia is an aphasia quotient of 93.8). POI 2’s performance was therefore not distinguishable from the performance of neurotypical controls. Her KBIT scores were 115 (84th percentile) on the verbal composite assessment (across the two subtasks), 109 (73rd percentile) on the non-verbal assessment, and 117 (79th percentile) overall composite assessment. She answered 50 of the 52 questions correct on the Pyramids and Palm Trees task.

#### 1.3.3 POI 3

POI 3 performed within normal range on all language and general cognitive tasks that were administered to her. She got 87.5% correct on PPVT, 95% correct on TROG. WAB-R was not administered. POI 3’s performance was therefore not distinguishable from the performance of neurotypical controls. Her KBIT scores were 145 (99.9th percentile) on the verbal composite assessment (across the two subtasks), 125 (95^th^ percentile) on the non-verbal assessment, and 139 (99.5th percentile) overall composite assessment. Pyramids and Palm Trees was not administered.

**Table S1:**
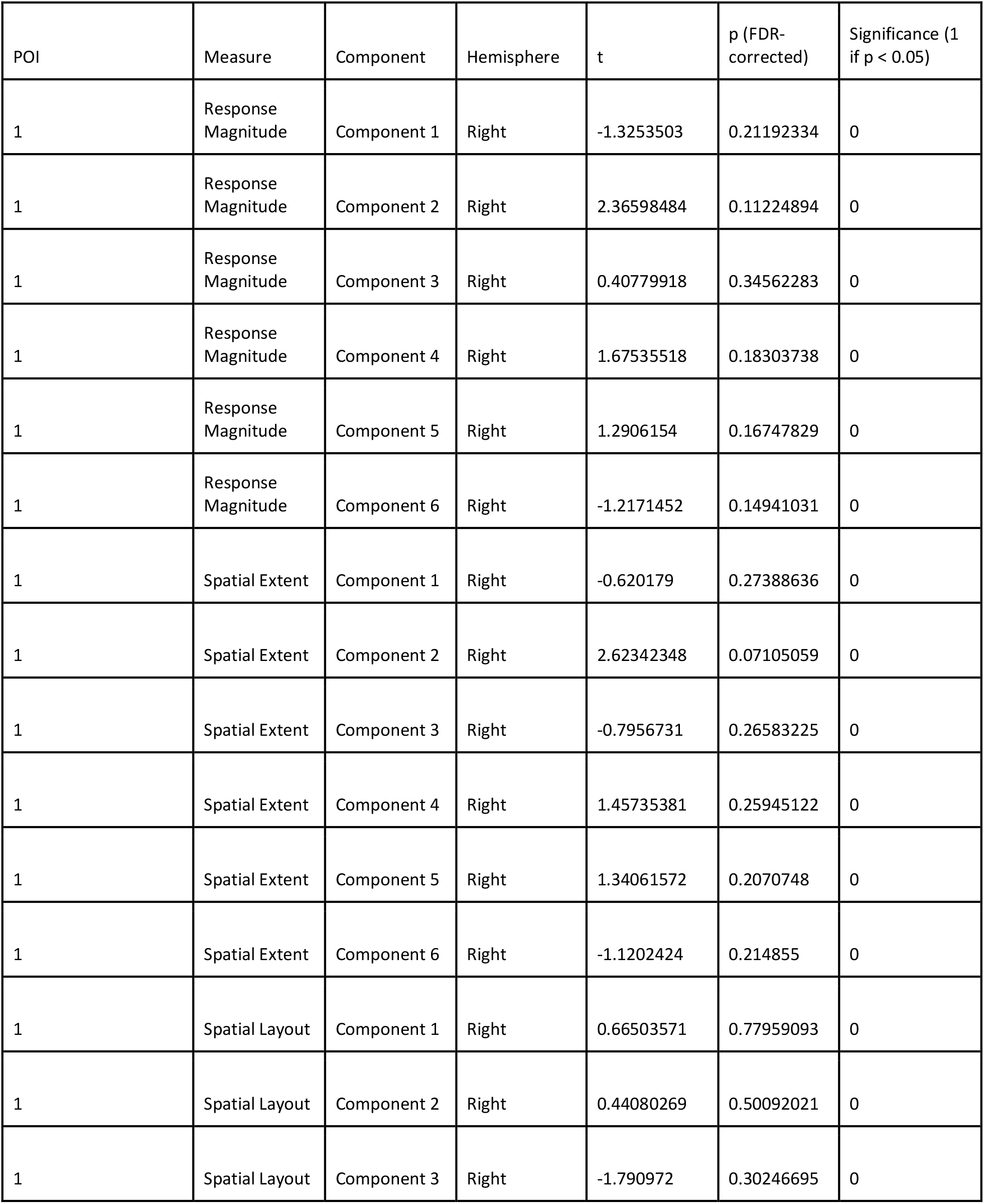
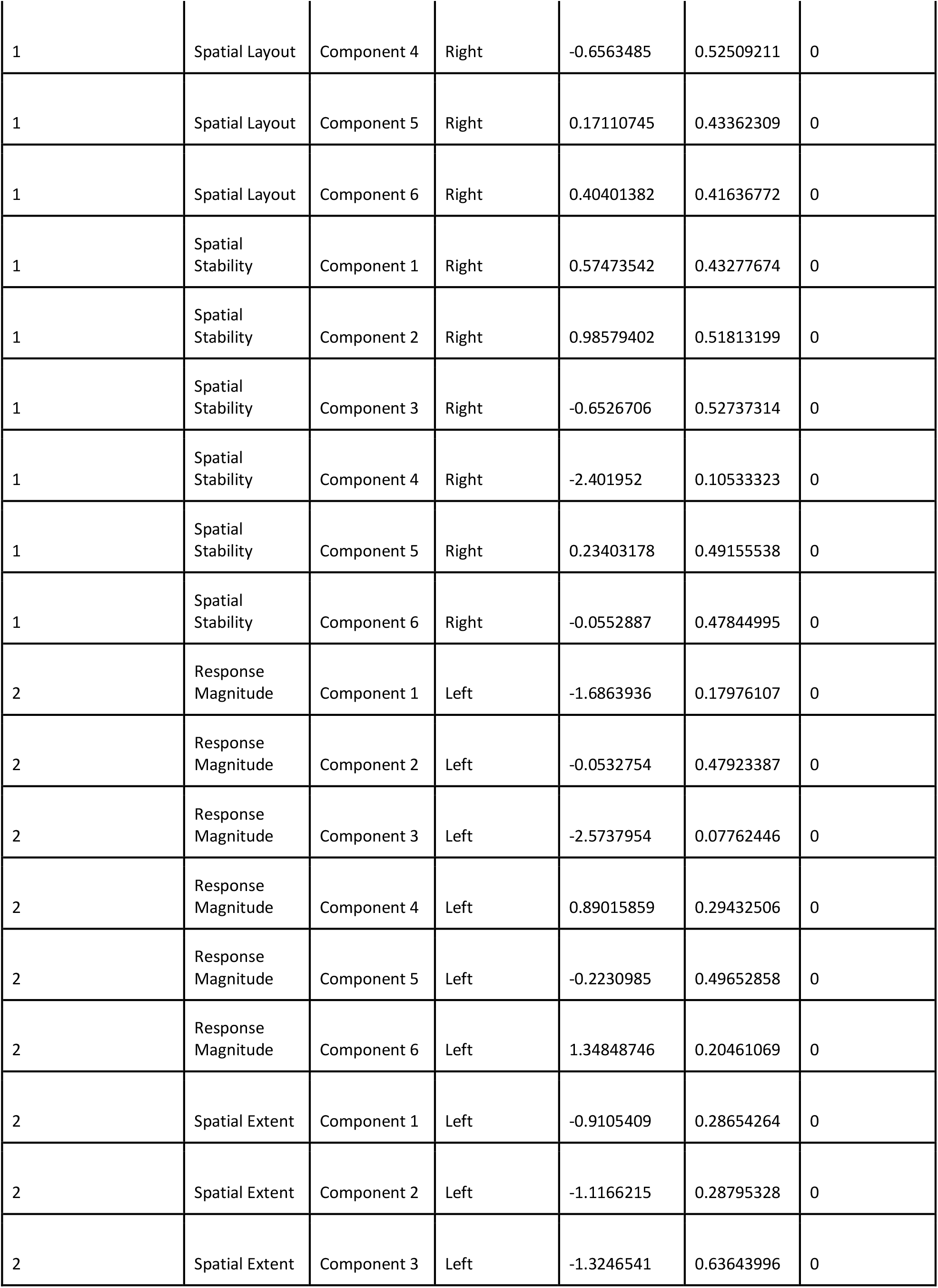
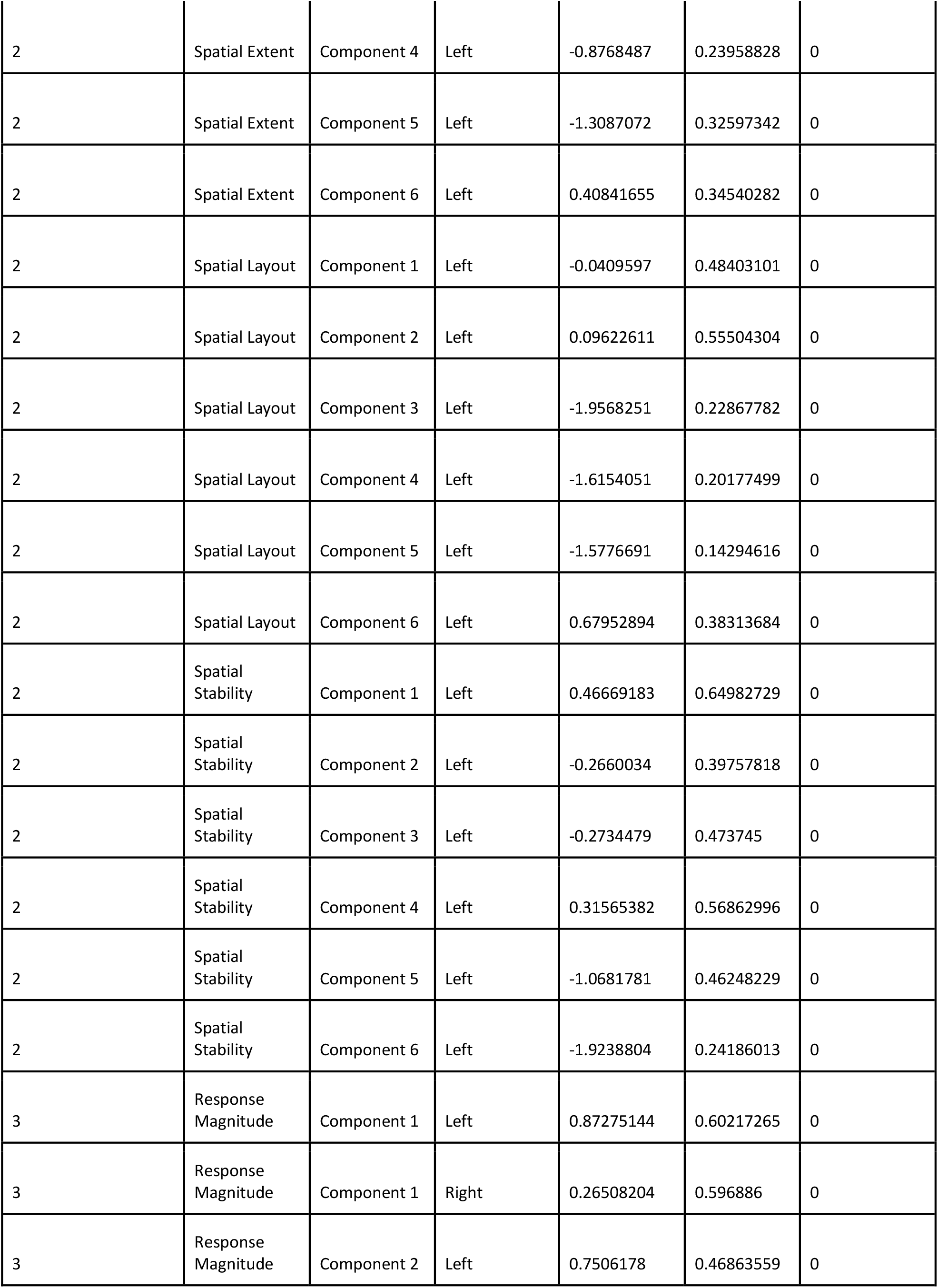
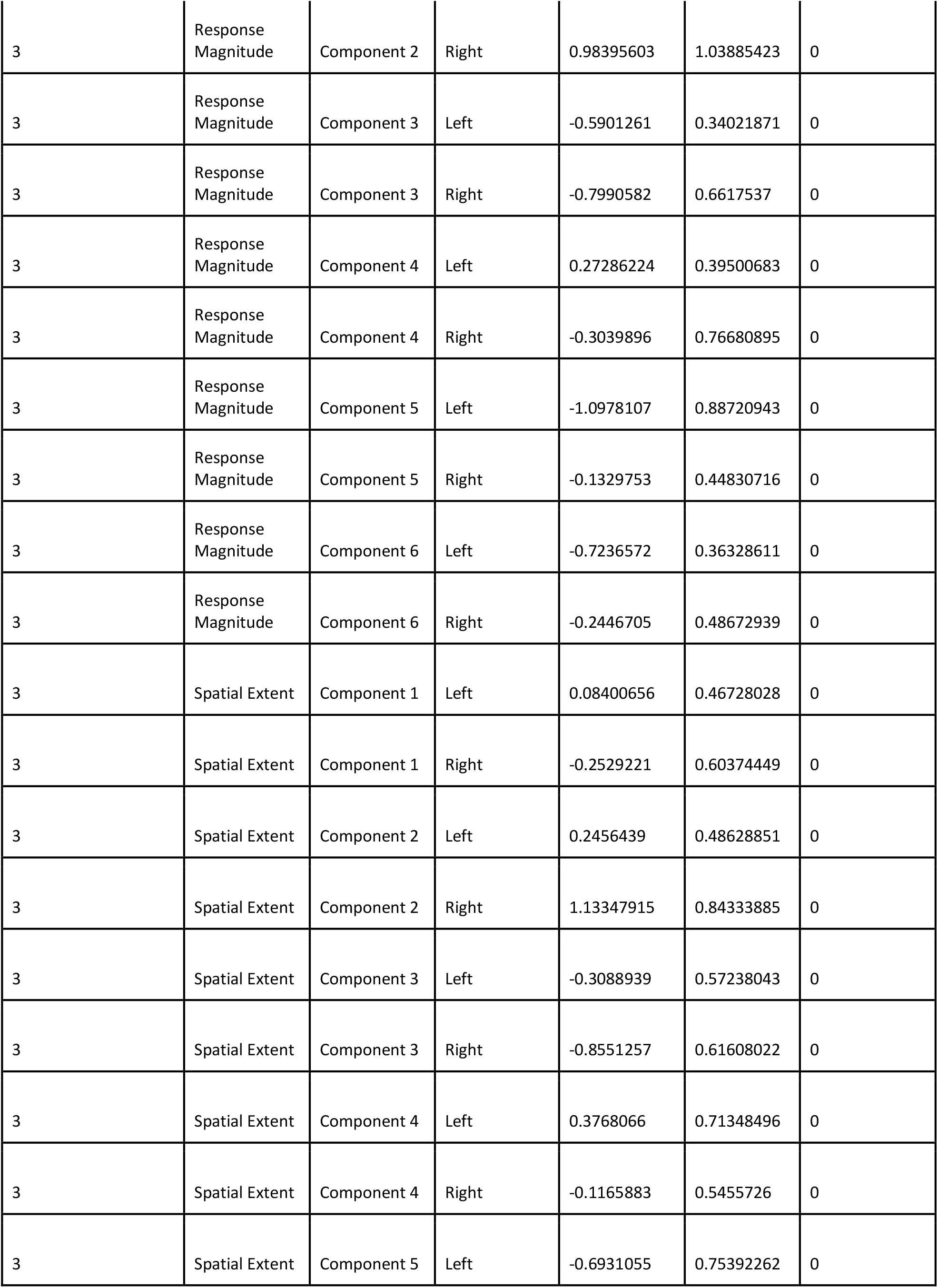
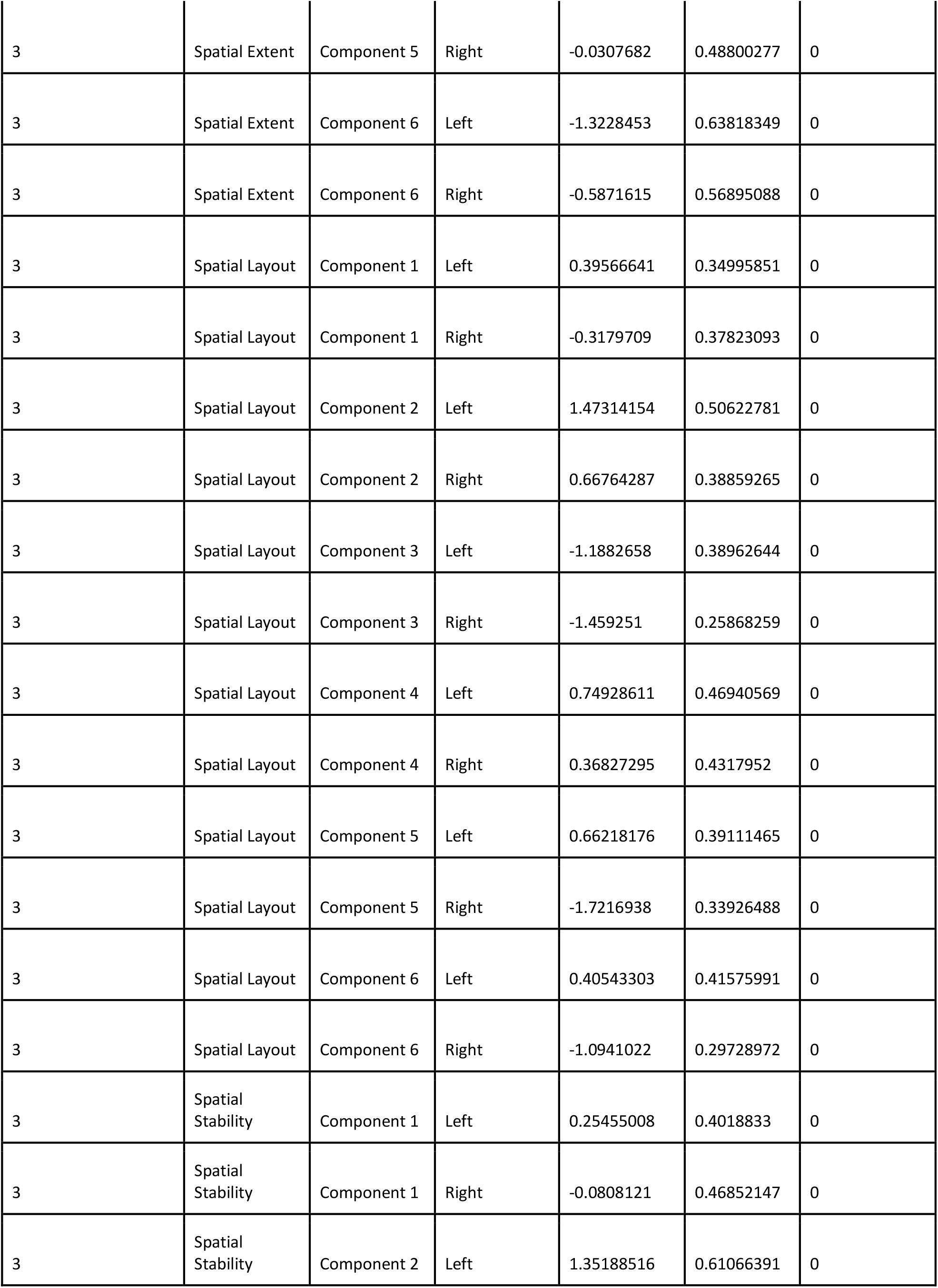
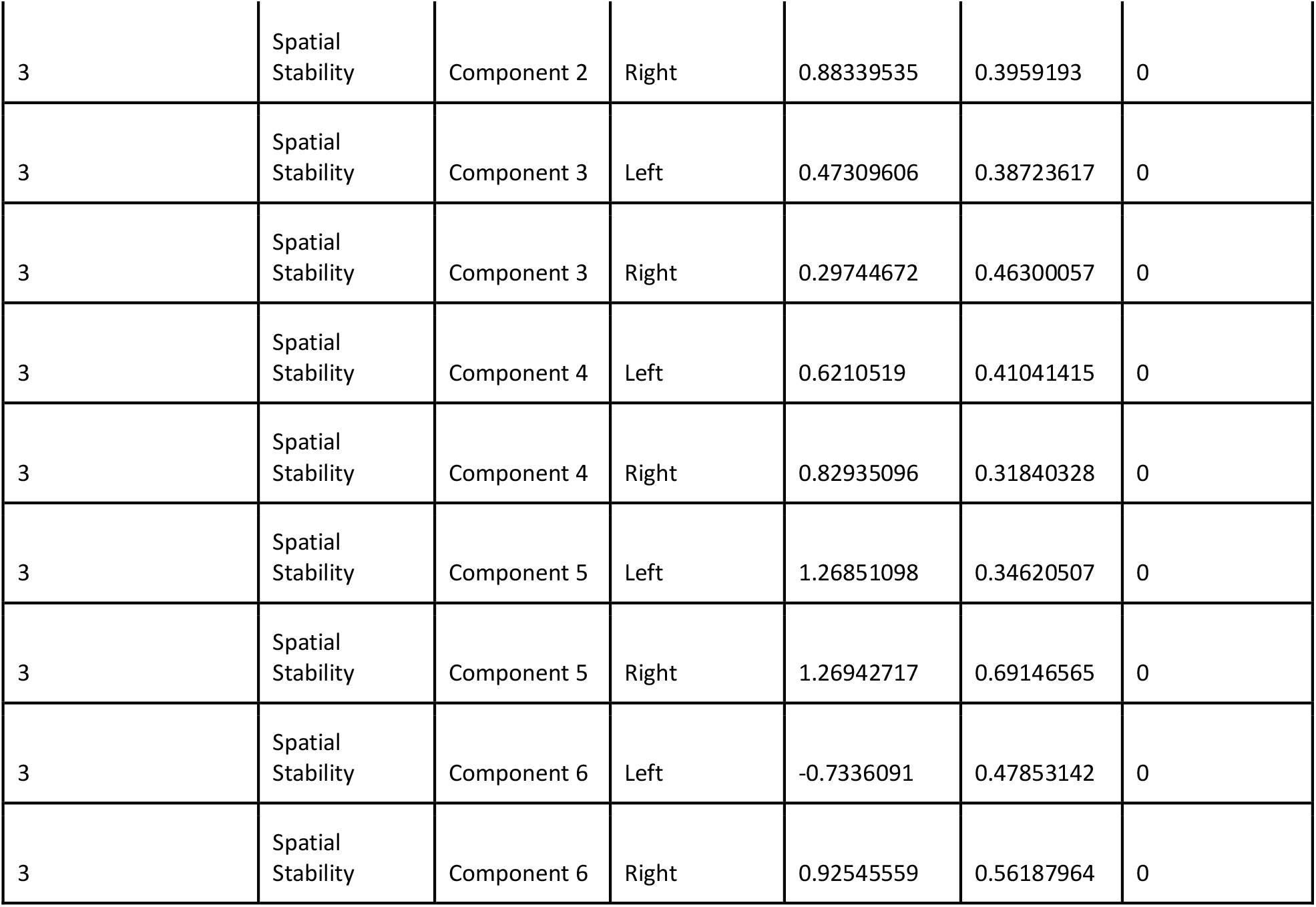
Compiled Crawford analysis results and statistics For each measure and component, the values of each POI (in each hemisphere for POI 3), were compared to the values of the 12 neurotypical controls using a Crawford test (Crawford & Howell, 1998, Methods) and the p-values were FDR corrected for the 6 components.

| POI | Measure | Component | Hemisphere | t | p (FDR-corrected) | Significance (1 if $p < 0.05$) |
| --- | --- | --- | --- | --- | --- | --- |
| 1 | Response Magnitude | Component 1 | Right | -1.3253503 | 0.21192334 | 0 |
| 1 | Response Magnitude | Component 2 | Right | 2.36598484 | 0.11224894 | 0 |
| 1 | Response Magnitude | Component 3 | Right | 0.40779918 | 0.34562283 | 0 |
| 1 | Response Magnitude | Component 4 | Right | 1.67535518 | 0.18303738 | 0 |
| 1 | Response Magnitude | Component 5 | Right | 1.2906154 | 0.16747829 | 0 |
| 1 | Response Magnitude | Component 6 | Right | -1.2171452 | 0.14941031 | 0 |
| 1 | Spatial Extent | Component 1 | Right | -0.620179 | 0.27388636 | 0 |
| 1 | Spatial Extent | Component 2 | Right | 2.62342348 | 0.07105059 | 0 |
| 1 | Spatial Extent | Component 3 | Right | -0.7956731 | 0.26583225 | 0 |
| 1 | Spatial Extent | Component 4 | Right | 1.45735381 | 0.25945122 | 0 |
| 1 | Spatial Extent | Component 5 | Right | 1.34061572 | 0.2070748 | 0 |
| 1 | Spatial Extent | Component 6 | Right | -1.1202424 | 0.214855 | 0 |
| 1 | Spatial Layout | Component 1 | Right | 0.66503571 | 0.77959093 | 0 |
| 1 | Spatial Layout | Component 2 | Right | 0.44080269 | 0.50092021 | 0 |
| 1 | Spatial Layout | Component 3 | Right | -1.790972 | 0.30246695 | 0 |
| 1 | Spatial Layout | Component 4 | Right | -0.6563485 | 0.52509211 | 0 |
| 1 | Spatial Layout | Component 5 | Right | 0.17110745 | 0.43362309 | 0 |
| 1 | Spatial Layout | Component 6 | Right | 0.40401382 | 0.41636772 | 0 |
| 1 | Spatial Stability | Component 1 | Right | 0.57473542 | 0.43277674 | 0 |
| 1 | Spatial Stability | Component 2 | Right | 0.98579402 | 0.51813199 | 0 |
| 1 | Spatial Stability | Component 3 | Right | -0.6526706 | 0.52737314 | 0 |
| 1 | Spatial Stability | Component 4 | Right | -2.401952 | 0.10533323 | 0 |
| 1 | Spatial Stability | Component 5 | Right | 0.23403178 | 0.49155538 | 0 |
| 1 | Spatial Stability | Component 6 | Right | -0.0552887 | 0.47844995 | 0 |
| 2 | Response Magnitude | Component 1 | Left | -1.6863936 | 0.17976107 | 0 |
| 2 | Response Magnitude | Component 2 | Left | -0.0532754 | 0.47923387 | 0 |
| 2 | Response Magnitude | Component 3 | Left | -2.5737954 | 0.07762446 | 0 |
| 2 | Response Magnitude | Component 4 | Left | 0.89015859 | 0.29432506 | 0 |
| 2 | Response Magnitude | Component 5 | Left | -0.2230985 | 0.49652858 | 0 |
| 2 | Response Magnitude | Component 6 | Left | 1.34848746 | 0.20461069 | 0 |
| 2 | Spatial Extent | Component 1 | Left | -0.9105409 | 0.28654264 | 0 |
| 2 | Spatial Extent | Component 2 | Left | -1.1166215 | 0.28795328 | 0 |
| 2 | Spatial Extent | Component 3 | Left | -1.3246541 | 0.63643996 | 0 |
| 2 | Spatial Extent | Component 4 | Left | -0.8768487 | 0.23958828 | 0 |
| 2 | Spatial Extent | Component 5 | Left | -1.3087072 | 0.32597342 | 0 |
| 2 | Spatial Extent | Component 6 | Left | 0.40841655 | 0.34540282 | 0 |
| 2 | Spatial Layout | Component 1 | Left | -0.0409597 | 0.48403101 | 0 |
| 2 | Spatial Layout | Component 2 | Left | 0.09622611 | 0.55504304 | 0 |
| 2 | Spatial Layout | Component 3 | Left | -1.9568251 | 0.22867782 | 0 |
| 2 | Spatial Layout | Component 4 | Left | -1.6154051 | 0.20177499 | 0 |
| 2 | Spatial Layout | Component 5 | Left | -1.5776691 | 0.14294616 | 0 |
| 2 | Spatial Layout | Component 6 | Left | 0.67952894 | 0.38313684 | 0 |
| 2 | Spatial Stability | Component 1 | Left | 0.46669183 | 0.64982729 | 0 |
| 2 | Spatial Stability | Component 2 | Left | -0.2660034 | 0.39757818 | 0 |
| 2 | Spatial Stability | Component 3 | Left | -0.2734479 | 0.473745 | 0 |
| 2 | Spatial Stability | Component 4 | Left | 0.31565382 | 0.56862996 | 0 |
| 2 | Spatial Stability | Component 5 | Left | -1.0681781 | 0.46248229 | 0 |
| 2 | Spatial Stability | Component 6 | Left | -1.9238804 | 0.24186013 | 0 |
| 3 | Response Magnitude | Component 1 | Left | 0.87275144 | 0.60217265 | 0 |
| 3 | Response Magnitude | Component 1 | Right | 0.26508204 | 0.596886 | 0 |
| 3 | Response Magnitude | Component 2 | Left | 0.7506178 | 0.46863559 | 0 |
| 3 | Response Magnitude | Component 2 | Right | 0.98395603 | 1.03885423 | 0 |
| 3 | Response Magnitude | Component 3 | Left | -0.5901261 | 0.34021871 | 0 |
| 3 | Response Magnitude | Component 3 | Right | -0.7990582 | 0.6617537 | 0 |
| 3 | Response Magnitude | Component 4 | Left | 0.27286224 | 0.39500683 | 0 |
| 3 | Response Magnitude | Component 4 | Right | -0.3039896 | 0.76680895 | 0 |
| 3 | Response Magnitude | Component 5 | Left | -1.0978107 | 0.88720943 | 0 |
| 3 | Response Magnitude | Component 5 | Right | -0.1329753 | 0.44830716 | 0 |
| 3 | Response Magnitude | Component 6 | Left | -0.7236572 | 0.36328611 | 0 |
| 3 | Response Magnitude | Component 6 | Right | -0.2446705 | 0.48672939 | 0 |
| 3 | Spatial Extent | Component 1 | Left | 0.08400656 | 0.46728028 | 0 |
| 3 | Spatial Extent | Component 1 | Right | -0.2529221 | 0.60374449 | 0 |
| 3 | Spatial Extent | Component 2 | Left | 0.2456439 | 0.48628851 | 0 |
| 3 | Spatial Extent | Component 2 | Right | 1.13347915 | 0.84333885 | 0 |
| 3 | Spatial Extent | Component 3 | Left | -0.3088939 | 0.57238043 | 0 |
| 3 | Spatial Extent | Component 3 | Right | -0.8551257 | 0.61608022 | 0 |
| 3 | Spatial Extent | Component 4 | Left | 0.3768066 | 0.71348496 | 0 |
| 3 | Spatial Extent | Component 4 | Right | -0.1165883 | 0.5455726 | 0 |
| 3 | Spatial Extent | Component 5 | Left | -0.6931055 | 0.75392262 | 0 |
| 3 | Spatial Extent | Component 5 | Right | -0.0307682 | 0.48800277 | 0 |
| 3 | Spatial Extent | Component 6 | Left | -1.3228453 | 0.63818349 | 0 |
| 3 | Spatial Extent | Component 6 | Right | -0.5871615 | 0.56895088 | 0 |
| 3 | Spatial Layout | Component 1 | Left | 0.39566641 | 0.34995851 | 0 |
| 3 | Spatial Layout | Component 1 | Right | -0.3179709 | 0.37823093 | 0 |
| 3 | Spatial Layout | Component 2 | Left | 1.47314154 | 0.50622781 | 0 |
| 3 | Spatial Layout | Component 2 | Right | 0.66764287 | 0.38859265 | 0 |
| 3 | Spatial Layout | Component 3 | Left | -1.1882658 | 0.38962644 | 0 |
| 3 | Spatial Layout | Component 3 | Right | -1.459251 | 0.25868259 | 0 |
| 3 | Spatial Layout | Component 4 | Left | 0.74928611 | 0.46940569 | 0 |
| 3 | Spatial Layout | Component 4 | Right | 0.36827295 | 0.4317952 | 0 |
| 3 | Spatial Layout | Component 5 | Left | 0.66218176 | 0.39111465 | 0 |
| 3 | Spatial Layout | Component 5 | Right | -1.7216938 | 0.33926488 | 0 |
| 3 | Spatial Layout | Component 6 | Left | 0.40543303 | 0.41575991 | 0 |
| 3 | Spatial Layout | Component 6 | Right | -1.0941022 | 0.29728972 | 0 |
| 3 | Spatial Stability | Component 1 | Left | 0.25455008 | 0.4018833 | 0 |
| 3 | Spatial Stability | Component 1 | Right | -0.0808121 | 0.46852147 | 0 |
| 3 | Spatial Stability | Component 2 | Left | 1.35188516 | 0.61066391 | 0 |
| 3 | Spatial Stability | Component 2 | Right | 0.88339535 | 0.3959193 | 0 |
| 3 | Spatial Stability | Component 3 | Left | 0.47309606 | 0.38723617 | 0 |
| 3 | Spatial Stability | Component 3 | Right | 0.29744672 | 0.46300057 | 0 |
| 3 | Spatial Stability | Component 4 | Left | 0.6210519 | 0.41041415 | 0 |
| 3 | Spatial Stability | Component 4 | Right | 0.82935096 | 0.31840328 | 0 |
| 3 | Spatial Stability | Component 5 | Left | 1.26851098 | 0.34620507 | 0 |
| 3 | Spatial Stability | Component 5 | Right | 1.26942717 | 0.69146565 | 0 |
| 3 | Spatial Stability | Component 6 | Left | -0.7336091 | 0.47853142 | 0 |
| 3 | Spatial Stability | Component 6 | Right | 0.92545559 | 0.56187964 | 0 |

**Table S2:** Correlation analyses results and statistics. Spatial Layout: Correlations between the voxel weights, averaged within each of 15 anatomical auditory cortex parcels (Glasser et al. 2016, Boebinger et al. 2021, Methods), of each POI and the average of the 12 neurotypical controls. Spatial Stability: Correlations between the voxel weights, averaged within each of 15 anatomical auditory cortex parcels (Glasser et al. 2016, Boebinger et al. 2021, Methods), of each POI in odd and even runs.

| POI | Measure | Component | Hemisphere | r* | p (FDR-corrected) | Significance (1 if $p < 0.05$) |
| --- | --- | --- | --- | --- | --- | --- |
| 1 | Spatial Layout | Component 1 | Right | 0.96188912 | 6.43E-08 | 1 |
| 1 | Spatial Layout | Component 2 | Right | 0.77677873 | 0.00131604 | 1 |
| 1 | Spatial Layout | Component 3 | Right | 0.15775647 | 0.57444426 | 0 |
| 1 | Spatial Layout | Component 4 | Right | 0.5140605 | 0.05994799 | 0 |
| 1 | Spatial Layout | Component 5 | Right | 0.85813313 | 0.0001278 | 1 |
| 1 | Spatial Layout | Component 6 | Right | 0.77589124 | 0.00101047 | 1 |
| 1 | Spatial Stability | Component 1 | Right | 0.87460204 | 5.97E-05 | 1 |
| 1 | Spatial Stability | Component 2 | Right | 0.90712025 | 1.84E-05 | 1 |
| 1 | Spatial Stability | Component 3 | Right | 0.44648617 | 0.14286803 | 0 |
| 1 | Spatial Stability | Component 4 | Right | 0.10402454 | 0.71217487 | 0 |
| 1 | Spatial Stability | Component 5 | Right | 0.8170039 | 0.00040143 | 1 |
| 1 | Spatial Stability | Component 6 | Right | 0.41374508 | 0.15030945 | 0 |
| 2 | Spatial Layout | Component 1 | Left | 0.76965633 | 0.00475294 | 1 |
| 2 | Spatial Layout | Component 2 | Left | 0.70621793 | 0.00976063 | 1 |
| 2 | Spatial Layout | Component 3 | Left | -0.3092621 | 0.26201844 | 0 |
| 2 | Spatial Layout | Component 4 | Left | 0.39572311 | 0.17312803 | 0 |
| 2 | Spatial Layout | Component 5 | Left | 0.54512142 | 0.05339118 | 0 |
| 2 | Spatial Layout | Component 6 | Left | 0.694366 | 0.00814967 | 1 |
| 2 | Spatial Stability | Component 1 | Left | 0.76690293 | 0.00509766 | 1 |
| 2 | Spatial Stability | Component 2 | Left | 0.64047771 | 0.02020553 | 1 |
| 2 | Spatial Stability | Component 3 | Left | 0.30857808 | 0.2631304 | 0 |
| 2 | Spatial Stability | Component 4 | Left | 0.61693268 | 0.02143526 | 1 |
| 2 | Spatial Stability | Component 5 | Left | 0.69254618 | 0.012643 | 1 |
| 2 | Spatial Stability | Component 6 | Left | -0.3246061 | 0.28539311 | 0 |
| 3 | Spatial Layout | Component 1 | Left | 0.89819817 | 1.63E-05 | 1 |
| 3 | Spatial Layout | Component 1 | Right | 0.65120338 | 0.01282057 | 1 |
| 3 | Spatial Layout | Component 2 | Left | 0.90153177 | 2.65E-05 | 1 |
| 3 | Spatial Layout | Component 2 | Right | 0.84328695 | 0.00047011 | 1 |
| 3 | Spatial Layout | Component 3 | Left | -0.0263637 | 0.92569446 | 0 |
| 3 | Spatial Layout | Component 3 | Right | 0.22591976 | 0.41815651 | 0 |
| 3 | Spatial Layout | Component 4 | Left | 0.74884661 | 0.00197368 | 1 |
| 3 | Spatial Layout | Component 4 | Right | 0.76010659 | 0.00301798 | 1 |
| 3 | Spatial Layout | Component 5 | Left | 0.85939839 | 8.06E-05 | 1 |
| 3 | Spatial Layout | Component 5 | Right | 0.70994109 | 0.00604968 | 1 |
| 3 | Spatial Layout | Component 6 | Left | 0.61432339 | 0.01779133 | 1 |
| 3 | Spatial Layout | Component 6 | Right | 0.58917141 | 0.02499031 | 1 |
| 3 | Spatial Stability | Component 1 | Left | 0.66392271 | 0.01042871 | 1 |
| 3 | Spatial Stability | Component 1 | Right | 0.61432071 | 0.01482666 | 1 |
| 3 | Spatial Stability | Component 2 | Left | 0.93487332 | 9.81E-07 | 1 |
| 3 | Spatial Stability | Component 2 | Right | 0.88446904 | 3.59E-05 | 1 |
| 3 | Spatial Stability | Component 3 | Left | 0.65290382 | 0.00998244 | 1 |
| 3 | Spatial Stability | Component 3 | Right | 0.65236129 | 0.01006924 | 1 |
| 3 | Spatial Stability | Component 4 | Left | 0.75295503 | 0.00238983 | 1 |
| 3 | Spatial Stability | Component 4 | Right | 0.78774784 | 0.00097598 | 1 |
| 3 | Spatial Stability | Component 5 | Left | 0.97745385 | 2.20E-09 | 1 |
| 3 | Spatial Stability | Component 5 | Right | 0.95713151 | 1.37E-07 | 1 |
| 3 | Spatial Stability | Component 6 | Left | 0.14317936 | 0.6107106 | 0 |
| 3 | Spatial Stability | Component 6 | Right | 0.68416597 | 0.00735969 | 1 |

\* These r values are marked as horizontal lines in the violin plots of Figure 4C and D (C-Spatial Layout, D-Spatial Stability).

## Notes

https://osf.io/qrx5n/

### Competing Interest Statement

The authors have declared no competing interest.

